# High-throughput and all-solution phase African Swine Fever Virus (ASFV) detection using CRISPR-Cas12a and fluorescence based point-of-care system

**DOI:** 10.1101/788489

**Authors:** Qian He, Dongmei Yu, Mengdi Bao, Grant Korensky, Juhong Chen, Mingyeong Shin, Juwon Kim, Myeongkee Park, Peiwu Qin, Ke Du

## Abstract

Here we report the development of a high throughput, all-solution phase, and isothermal detection system to detect African Swine Fever Virus (ASFV). CRISPR-Cas12a programmed with a CRISPR RNA (crRNA) is used to detect ASFV target DNA. Upon ASFV DNA binding, the Cas12a/crRNA/ASFV DNA complex becomes activated and degrades a fluorescent single stranded DNA (ssDNA) reporter present in the assay. We combine this powerful CRISPR-Cas assay with fluorescence-based point-of-care (POC) system we developed for rapid and accurate virus detection. Without nucleic acid amplification, a detection limit of 1 pM is achieved within 2 hrs. In addition, the ternary Cas12a/crRNA/ASFV DNA complex is highly stable at physiological temperature and continues to cleave the ssDNA reporter even after 24 hrs of incubation, resulting in an improvement of the detection limit to 100 fM. We show that this system is very specific and can differentiate nucleic acid targets with closely matched sequences. The high sensitivity and selectivity of our system enables the detection of ASFV in femtomolar range. Importantly, this system features a disposable cartridge and a sensitive custom designed fluorometer, enabling compact, multiplexing, and simple ASFV detection, intended for low resource settings.

## 1. Introduction

African swine fever virus (ASFV) is a highly virulent pathogen that causes near 100% mortality rate in domestic pigs and wild boars (Galindo and Alonso, 2017; Sánchez-Cordón et al., 2018). Its high persistence in hosts, pork products, soft ticks, and swills has made it extremely difficult to control, and thus is becoming a global threat to food industry and international trade. In August 2018, the ASFV outbreak in Asia was reported as the biggest animal disease outbreak to date, killing over 500 million pigs worldwide and putting thousands of farmers out of business (https://www.pirbright.ac.uk/press-releases/2018/08/pirbright-scientists-help-combat-african-swine-fever-disease-no-vaccine). Although ASFV cannot infect humans, the impact of the disease in Asia has been causing significant economic losses and shows no signs to slowing down (https://www.theguardian.com/world/2019/sep/09/philippines-confirms-first-swine-fever-cases). Moreover, heparin, a polymer that is widely used as an anticoagulant for the treatment of heart attack and unstable angina, could face a global shortage because nearly 80% of crude heparin is derived from mucosal tissues of pigs (Vilanova et al., 2019).

Without effective treatment and vaccines, early and accurate diagnosis is an effective alternative way to rapidly control outbreaks in infected regions, followed by movement restrictions and stamping-out policies (Moennig, 2000). For regions that are free of the disease, a practically feasible ASFV detection method can enhance biosafety awareness and control. However, a reliable, inexpensive, highly sensitive, and high-throughput ASFV detection method that is suitable for point-of-care (POC) settings is still unexplored. The current gold standard for ASFV diagnostics in the laboratory is based on the traditional polymerase chain reaction (PCR), a viral genome detection technique based on nucleic acid marker amplification (Agüero et al., 2003; Luo et al., 2017). Even though highly sensitive, PCR is too complicated for field virus detection, requiring frozen storage of fragile enzymes, and relying on bulky, expensive instruments. All these shortcomings have prevented its wide-scale use in rural infected communities. Moreover, PCR is susceptible to cross-contamination in high-throughput screening, thus requiring skilled operators.

On the other hand, virus isolation based on viral replication in the susceptible primary cells has been used for ASFV diagnosis (Chapman et al., 2011; O’Donnell et al., 2016; Rowlands et al., 2008). However, the detection specificity of this virus isolation method is relatively poor and can result in misdiagnosis. The entire virus isolation detection process takes 48-72 hours, which is not ideal for the high-throughput and rapid detection in resource-limited settings. Enzyme-linked immunosorbent assay (ELISA), a technique of recognizing viral antigens, has also been widely used for ASFV detection (Cubillos et al., 2013; Giménez-Lirola et al., 2016). ELISA does not require expensive instruments and complicated amplification processes (Giménez-Lirola et al., 2016). However, ELISA has poor sensitivity and relies on fragile temperature-sensitive reagents (Gallardo et al., 2015). The reagents can easily be degraded in low resource settings, further decreasing its detection sensitivity. Thus, ELISA must be paired with other molecular diagnostic methods for pathogen detection in POC settings.

Clustered regularly interspaced short palindromic repeats (CRISPR) technology has emerged in recent years as a powerful tool to modify defective genes within living organisms for disease treatment (Canver et al., 2015; Dever et al., 2016; Knott et al., 2017; Schwank et al., 2013; Tambe et al., 2018) The recent discovery of CRISPR Cas12a and Cas13a, which possess the capability of recognizing a nucleic acid target, followed by the indiscriminate cleavage of nucleic acid reporters, has been used for rapid detection of pathogens (Chen et al., 2018; East-Seletsky et al., 2017, 2016; Gootenberg et al., 2017; Qin et al., 2019) Rather than using precisely controlled heating cycles in PCR, CRISPR-based detection can be operated at physiological temperature or even room temperature (Kundert et al., 2019; Malzahn et al., 2019; Qin et al., 2019), thus significantly simplifying the detection procedure. The non-specific cleavage by these CRISPR-Cas enzymes occur at a very high turnover rate, enabling accurate detection of low-concentration targets. In addition, the high robustness of CRISPR assay allows the reaction to happen in bodily fluids (Chen et al., 2018), avoiding complicated sample purification and preparation procedures, which is a key benefit for rapid and low-cost detection. We previously developed a fully automated microfluidic system with an integrated fluorescence sensing unit for all-solution phase Ebola detection with Cas13a (Qin et al., 2019). A detection limit of 5.45×10^7^ copies/mL was achieved without target amplification, thus establishing a powerful and quantitative POC system for pathogen detection.

Here, we present an advanced system for rapid and accurate detection of ASFV. Because ASFV possesses a double-stranded DNA genome, we developed a CRISPR Cas12a detection scheme by cleaving single-stranded DNA probes after ASFV target hybridization with a crRNA-programmed Cas12a. The small fluorescence sensing unit is aligned with an 80x disposable cartridge, thus matching the high throughput of commercialized PCR systems. We were able to distinguish closely matched target samples with a detection limit of ∼1 pM. Furthermore, we demonstrated the high stability of Cas12a system, extending the detection limit to 100 fM with longer incubation time. This compact detection system is automated, integrated, small, lightweight, and inexpensive, ready to be used for on-site ASFV detection or other DNA based pathogen detection.

## 2. Materials and Methods

### 2.1 Chemicals and reagents

ASFV-SY18, the strain corresponding to the 2018 ASFV outbreak in China (Zhou et al., 2018), was chosen as the target. B646L gene was selected because it is a highly conserved gene encoding vp72, an essential protein that is required for infection of cells. Synthetic double stranded DNA (dsDNA) oligonucleotides, including ASFV-DNA (A-DNA) and mutated ASFV-DNA (M-DNA) were purchased from Integrated DNA Technologies (IDT, Inc.). Both of these dsDNA oligonucleotides have two strands, one is target strand (TS) and the other is non-target strand (NTS). We designed a 5 base pair difference of A-DNA and M-DNA after the PAM region. CRISPR RNA (crRNA), single-stranded DNA (ssDNA) probes including DNaseAlert and a custom-designed reporter (F-Q), IDTE buffer, and nuclease free water were all purchased from Integrated DNA Technologies (IDT, Inc.). *Lachnospiraceae bacterium* Cas12a (LbCas12a) proteins at analytical grade were purchased from New England BioLabs, USA and used without any further purification. The sequences of all the oligonucleotides used in this work are shown in Table 1.

**Table 1.**
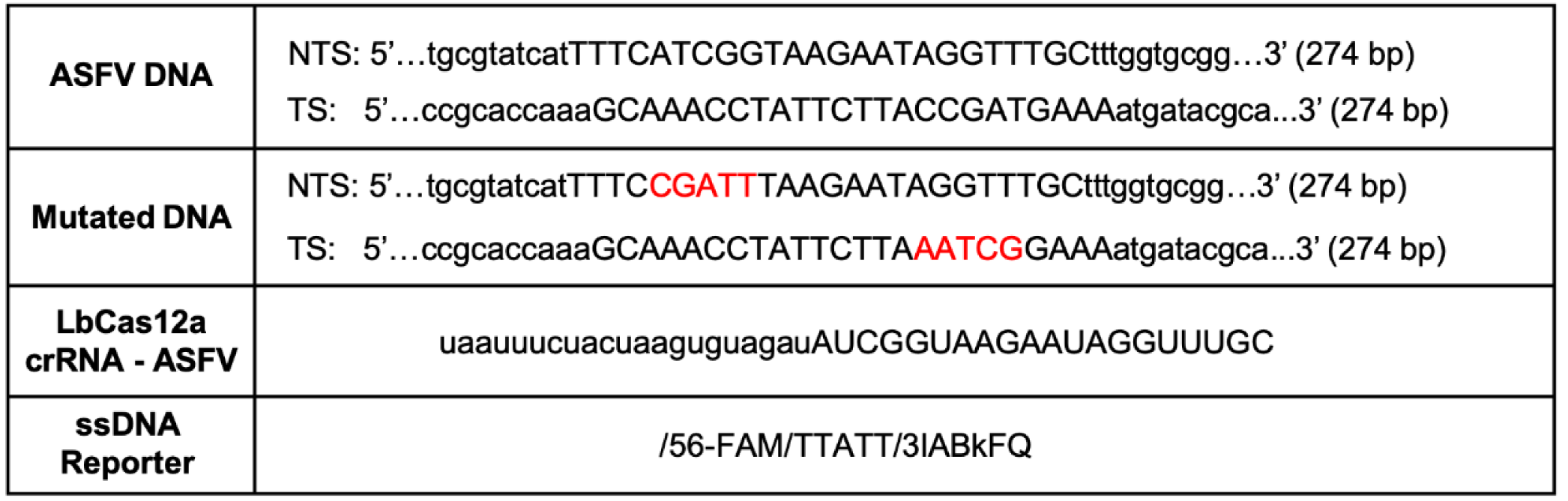
Sequence of DNA and crRNA.

### 2.2 Reaction assay

LbCas12a-crRNA complexes were preassembled by incubating 1 µM LbCas12a and 1.25 µM crRNA at room temperature for 10 min. DNA samples (A-DNA and M-DNA) with various concentrations were dissolved in 1× Binding Buffer (10 mM Tris-HCl, pH 7.5, 50 mM NaCl, 5 mM MgCl_2_, 0.1 mg/ml BSA), mixed with LbCas12a-crRNA complexes and 50 nM ssDNA reporter probe in a 20 μL reaction volume. In the assay, the final concentrations of LbCas12a and crRNA were 50 nM and 62.5 nM, respectively. The mixture was incubated at 37°C in a water bath for 2 to 24 hrs. The cleaved product was then added into a disposable cartridge and placed on our custom designed benchtop fluorometer for detection (**Fig. 1a**).

**Figure 1.**
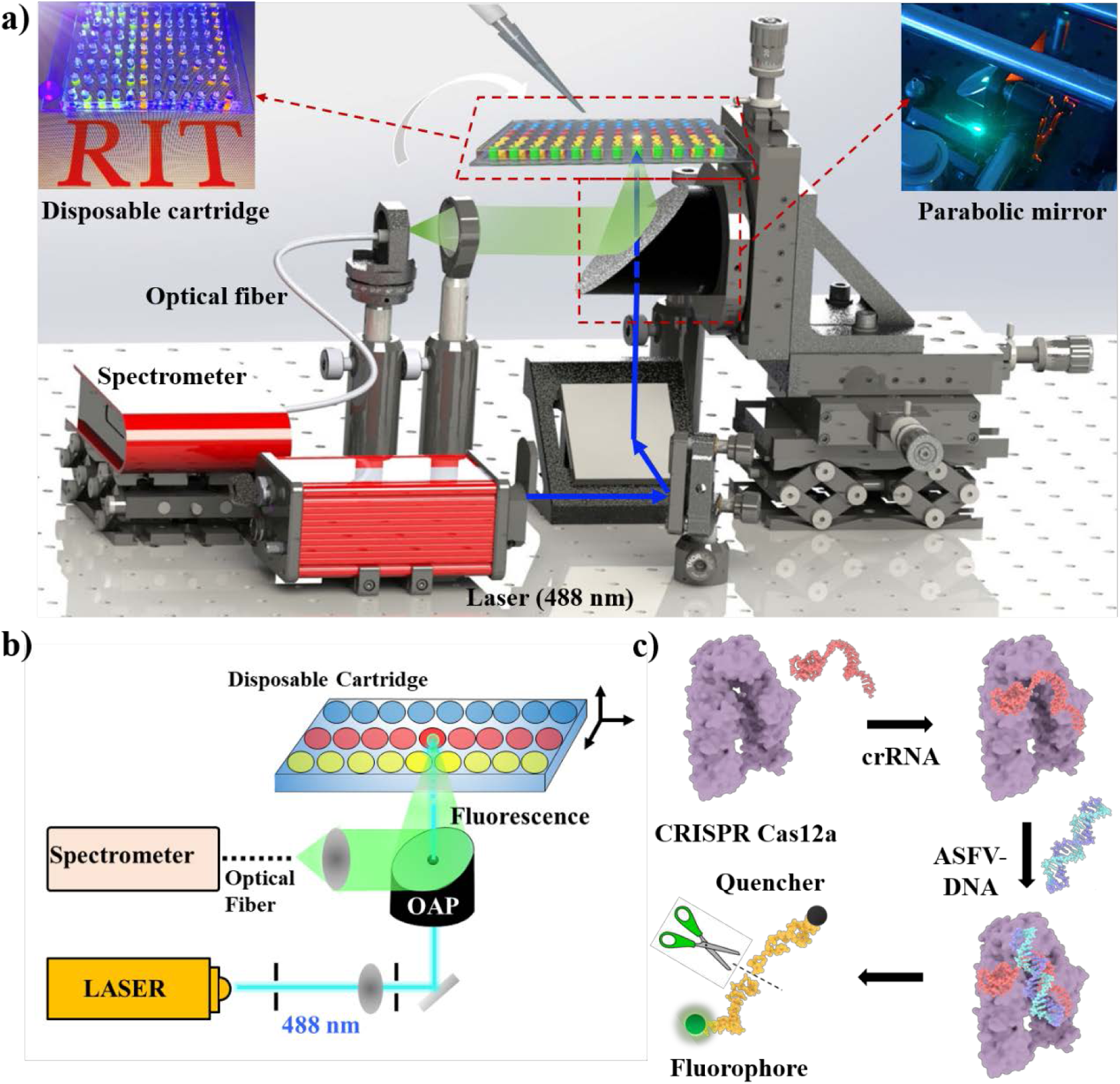
(a) Schematic of the POC system for rapid ASFV diagnostics. Insets: Photograph of the 80x disposable cartridge and the parabolic mirror setup. (b) Schematic of the fluorescence sensing unit with a 488 nm laser as an excitation source. (c) Schematic of the CRISPR Cas12a detection mechanism: CRISPR Cas12a binds with crRNA and ASFV-DNA and the Cas12a/crRNA/ASFV complex cleaves ssDNA probe for fluorescence sensing.

### 2.3 Components of measurement

The disposable cartridge was made by bonding a Polydimethylsiloxane (PDMS) slab on a glass substrate. Sylgard 184 (Dow Corning, USA) PDMS was prepared by mixing the base and curing agent with a ratio of 10:1, followed by baking on a hotplate at 80°C for 2 hrs. Eighty reservoirs with a diameter of 3.5 mm and a period of 11 mm were created using a biopsy punch. Both glass slide (L: 75 mm; W: 50 mm) and PDMS were treated with oxygen plasma treatment (Electro-Technic Products, USA) for 120 s and pressed together. Finally, the glass/PDMS sample was baked at 95°C for 2 hrs. The Si-O-Si bond permanently seals the two layers (Xiong et al., 2014).

The design of the integrated fluorescence sensing unit was reported before (Qin et al., 2019) and is shown in **Fig. 1b**. Briefly, the disposable cartridge was placed on the fluorometer and aligned by the three-axis translational stage. A continuous laser (488 nm) was used as an excitation source. To avoid photobleaching issues, the power was reduced to 3 mW. The fluorescence signal was collected using an off-axis parabolic (OAP) mirror and filtered by a 488 nm notch filter (Thorlabs, Inc.). The collected signal was focused into an optical fiber (M93L01, Thorlabs) and the spectra was recorded by a portable USB spectrometer (USB 2000+, Ocean Optics).

### 2.4 Fluorescence Detection

The detection mechanism is based on the non-specific ssDNA probe cleavage by LbCas12a as shown in **Fig. 1c**. To report the presence of target DNA, ssDNA probe linking a fluorophore and a quencher was used. LbCas12a first binds with crRNA. Then, LbCas12a-crRNA complex binds with A-DNA and the complex triggers the indiscriminate cleavage of ssDNA in the solution. The fluorophore on the ssDNA is released into the assay and detected by the fluorescence sensing unit.

## 3. Results

To validate the CRISPR Cas12a detection assay, two different ssDNA reporter probes with emission peaks at 556 nm and 520 nm were tested using a commercially available spectrofluorometer (JASCO FP-8500, USA). As presented in **Fig. 2a**, with an A-DNA concentration of 10 nM, the fluorescence intensity of 500 nM DNaseAlert is statistically significantly higher than 50 nM DNaseAlert, indicating that 50 nM DNaseAlert is not saturating the CRISPR-crRNA-ASFV DNA complex. The integrated signals of the fluorescence spectra are plotted towards DNaseAlert concentration (**Fig. 2b**). The integrated signal with 500 nM is ∼1,700 counts, which is approximately 3 times higher than that of 50 nM DNaseAlert. By using 500 nM of DNaseAlert, we then investigated the relationship between DNA concentration and fluorescent signal intensity by adding 0.1 nM, 1nM, and 10 nM A-DNA to reaction assay. The fluorescent signal linearly increases with A-DNA concentration from 0.1 nM to 10 nM (**Fig. 2d**). The linear correlation (Pearson’s R = 0.99) between fluorescent intensity and A-DNA concentration demonstrates that we are able to quantitatively measure different concentrations of A-DNA (**Fig. 2d**). Similarly, higher fluorescence signal is detected using higher concentration of the custom-designed reporter (F-Q). As shown in **Fig. 2e** and **Fig. 2f**, applying 500 nM F-Q generates ∼3 times more fluorescence signal than 50 nM F-Q. It also demonstrates excellent linearity (Pearson’s R = 0.98) by varying the target concentration from 0.1 nM to 10 nM.

**Figure 2.**
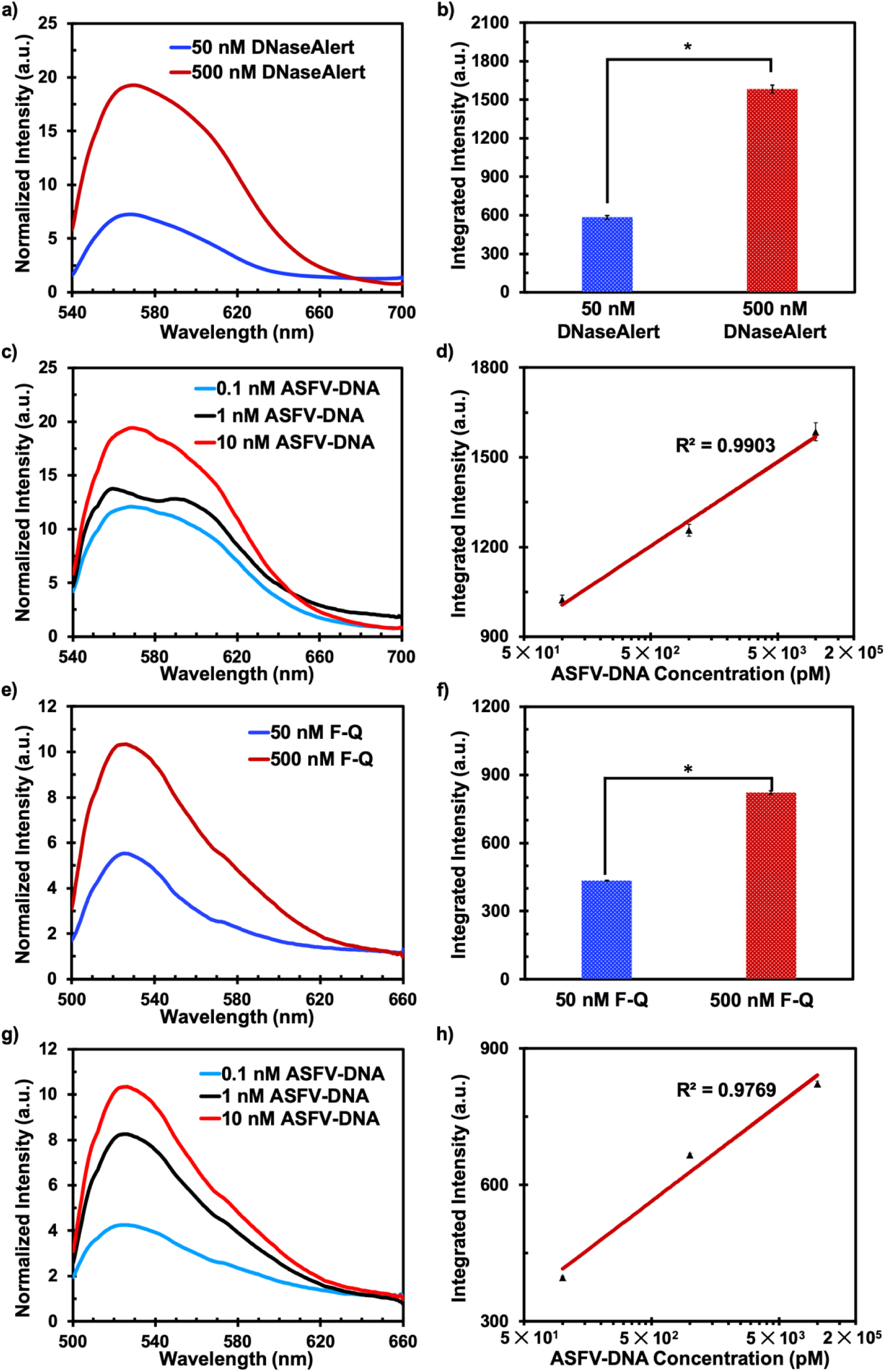
Detection of ASFV dsDNA by CRISP-Cas12a reaction system with two kinds of DNA reporters: DNaseAlert and ssDNA Fluorophore-Quencher (F-Q). (a) ASFV dsDNA detection with 50 nM and 500 nM DNaseAlert. (b) Integrated fluorescence signal of (a) with a wavelength from 540 nm to 700 nm. (c) Emission curve of ASFV dsDNA with various concentrations measured by 500 nM DNaseAlert. (d) Fluorescence intensity (540 nm-700 nm) versus ASFV dsDNA showing linear dependence. (e) ASFV dsDNA detection with 50 nM and 500 nM F-Q reporter. (f) Integrated fluorescence signal of (f) with a wavelength from 500 nm to 660 nm. (g) Emission curve of ASFV dsDNA with various concentrations by 500 nM F-Q. (h) Fluorescence intensity (500 nm-660 nm) versus ASFV dsDNA showing linear dependence.

After optimizing the detection assay, we integrated the CRISPR Cas12a assay with our sensitive benchtop POC system to determine the detection limit of ASFV. The incubation time was 2 hrs. As shown in **Fig. 3a**, the highest signal intensity is observed using 100 pM A-DNA (dashed purple), followed by 10 pM (dashed green) and 1 pM A-DNA (dashed black). On the other hand, the measured intensity of 100 pM M-DNA is only slightly higher than the background without any target DNA. And the measured signal of M-DNA with a concentration from 10 fM to 10 pM are all comparable to the background. We summarized the integrated signal versus target concentration of A-DNA and M-DNA and present it in **Fig. 3b**. The intensities of 1 pM, 10 pM, and 100 pM A-DNA are ∼300, ∼500, and ∼1,100 and are significantly higher than their related M-DNA and background.

**Figure 3.**
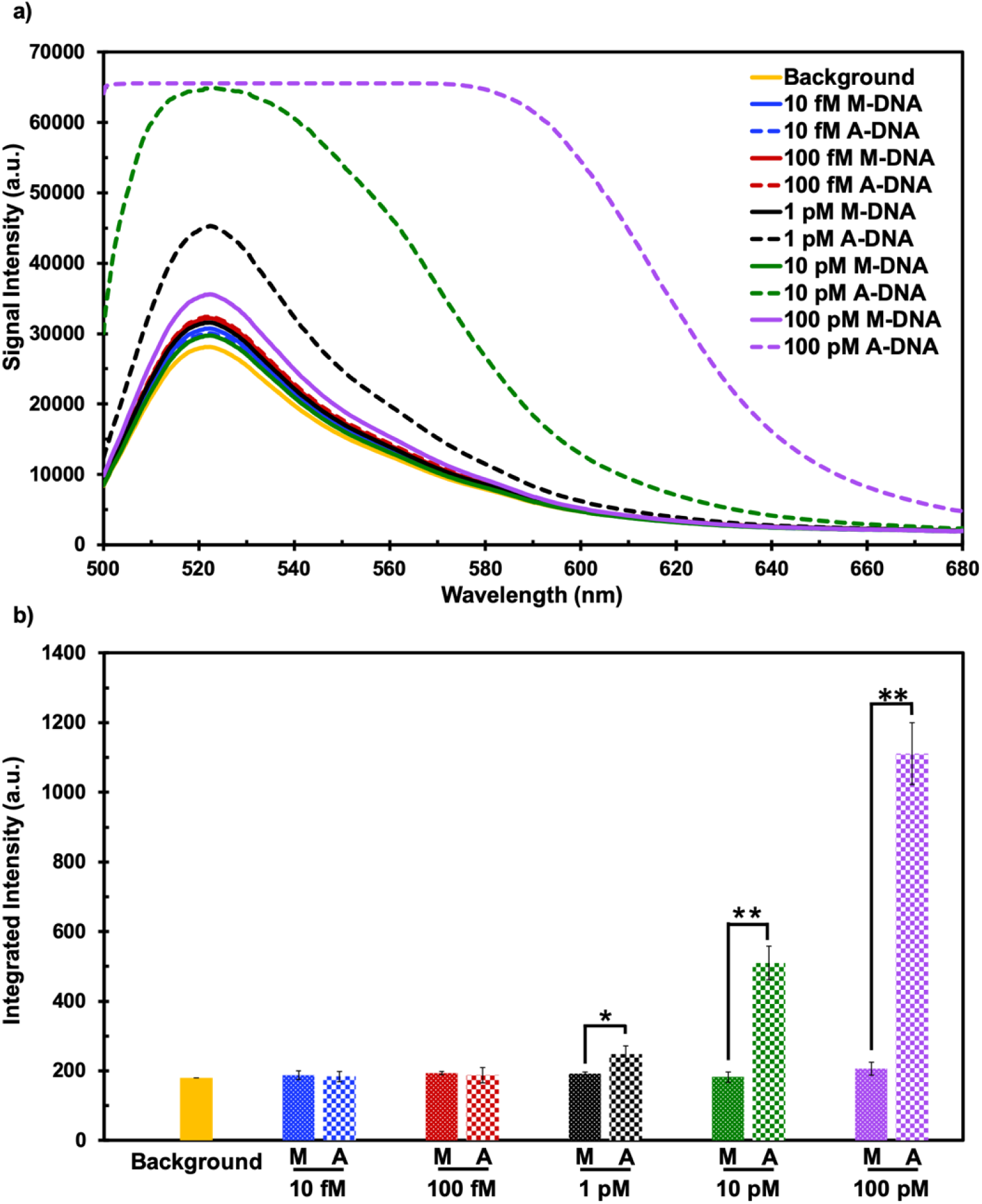
(a) Uncorrected emission curves of A-DNA (dashed line) and M-DNA (full line) with various concentrations detected by using custom designed POC system. (b) Integrated signal intensity of background (yellow), A-DNA (blocks), and M-DNA (dots) with various concentrations and an incubation time of 2 hrs.

We further explored the detection sensitivity versus incubation time with our POC system by extending the reaction time to 24 hrs (**Fig. 4**). After 8 hrs incubation, 1 pM A-DNA is easily detected while the intensity of other groups is comparable with the background. We found that the fluorescence intensity of 1 pM A-DNA and 100 fM A-DNA both increases with 16 hrs incubation time. After 24 hrs incubation, the fluorescence signal of 100 fM A-DNA is significantly higher than M-DNA (**: < 0.005), indicating that a detection limit of 100 fM A-DNA (5.7 × 10^7^ copies/mL) can be detected after 24 hrs incubation.

**Figure 4.**
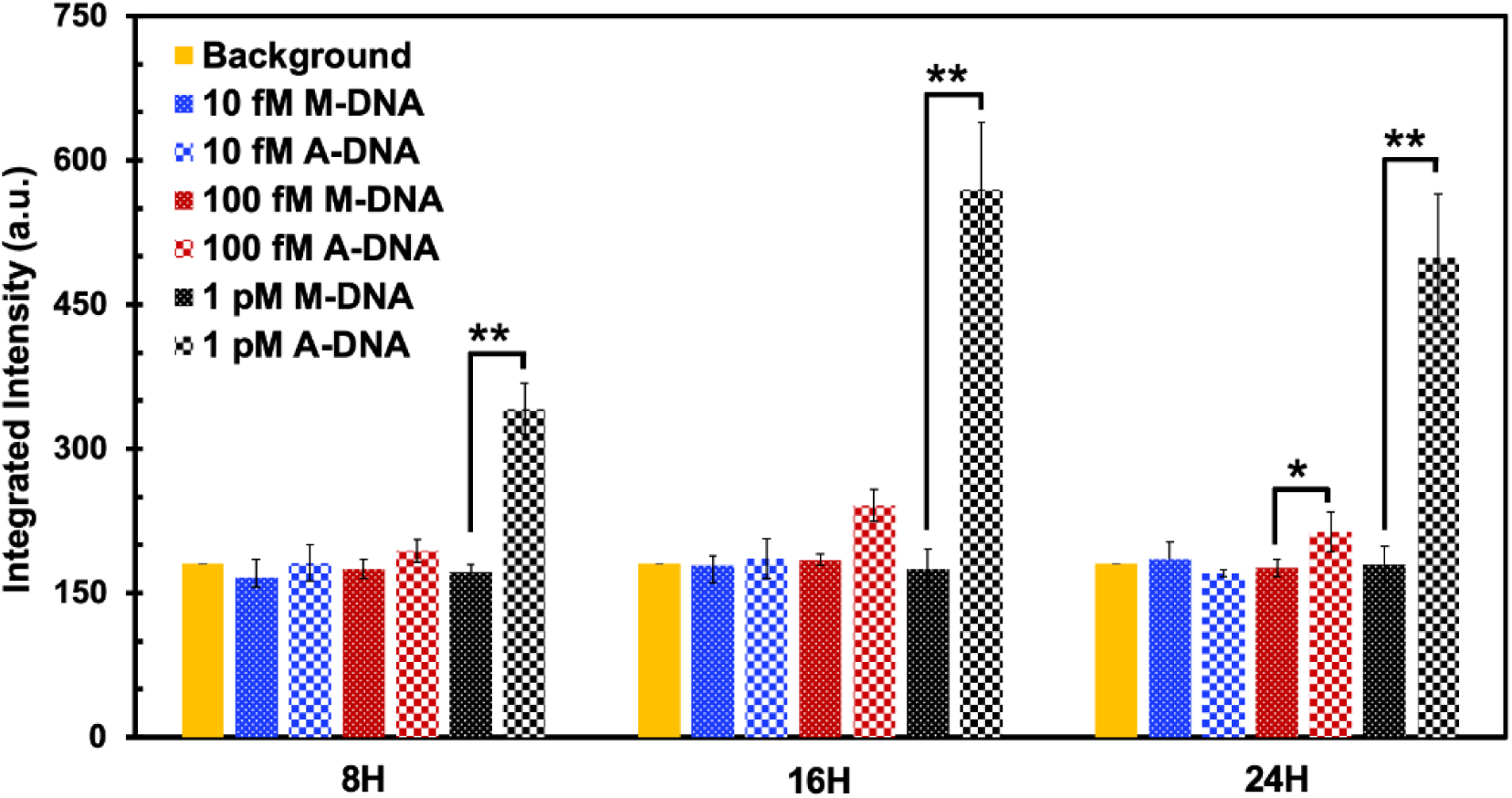
Integrated intensity of background (yellow), A-DNA (blocks), and M-DNA (dots) with a concentration ranging from 10 fM to 1 pM and an incubation time of 8 hrs, 16 hr, and 24 hrs. *: < 0.05, **: < 0.005.

## 4. Discussion

Our POC system using a portable laser and a mini spectrometer is ∼100 times more sensitive than commercially available spectrofluorometers and is much cheaper. Other techniques, such as fluorescence microscopy, have been introduced for viral particle imaging (Pereira et al., 2012; Wang et al., 2018). Fluorescence microscopes provide higher sensitivity and can trace single molecules. However, microscopy is bulky and very expensive: not suitable for rapid POC diagnosis. Furthermore, experts are needed to operate and analyze the results. Other detection methods such as electrochemical sensing and surface enhanced Raman microscopy have poor sensitivity and cannot be used to report the presence of pathogens when the viral load is low (Granger et al., 2016; Hsieh et al., 2015). These methods also rely on sophisticated nanofabrication to pattern uniform nanostructures, which could increase the costs of the sensing chip. Our approach combining a CRISPR Cas12a assay with a small fluorescence sensing unit is very easy to operate, doesn’t require expert operation, and shows excellent detection sensitivity at femtomolar level. It is achieved by simply mixing the CRISPR assay with pathological samples and then adding the mixture into a disposable cartridge for rapid measurement (Guk et al., 2017; Myhrvold et al., 2018). The detection is all in solution and does not require complicated solid phase extraction. In addition, the disposable cartridge is very simple to make and does not need microfabrication.

Using our approach, we report a detection limit of 100 fM (5.7 × 10^7^ copies/mL) for A-DNA without using nucleic acid amplification. Even though PCR shows higher sensitivity than our method, it requires expensive instrumentation and reagents for target amplification, which is not suitable for POC diagnostics (Su et al., 2015). The CRISPR Cas12a assay is isothermal and amplification-free, making it a possible substitute for PCR. Furthermore, the detection sensitivity of Cas12a assay can easily be extended to attomolar level by adding an isothermal recombinase polymerase amplification (RPA) (Chaijarasphong et al., 2019). Unlike PCR which requires thermal cycles, the Cas12a assay and RPA reaction are carried out at 37°C by mixing the reagents together without the need of precise temperature control.

LbCas12a protein is an RNA-guide enzyme that binds and cleaves ssDNA probes (Li et al., 2018). The Cas12a system is ideal for POC diagnostics towards DNA viruses due to its very simple process and high stability. Both Cas 12a and Cas13a have collateral activities and Cas12a can be activated by single stranded DNA target even without PAM (Chen et al., 2018). Compared to single stranded DNA target, the efficacy of non-specific ssDNA cleavage of Cas12a can be elevated by NTS from double stranded target DNA, thus providing excellent stabilization. In our research, we found that Cas12a is constantly activated for 24 hrs, due to the high stability of Cas12a-crRNA-target double stranded DNA complex.

Similar to Cas 9, Cas12a is also a DNA endonuclease. Unlike Cas9 which needs crRNA and tracrRNA, Cas12a is guided by only crRNA (Fonfara et al., 2016). Although the dual RNA guide fused tracrRNA and crRNA was reported with Cas9 (Jinek et al., 2012), the size of RNA molecule (117 bp) used for Cas9 detection is larger than the crRNA of Cas12a (41 bp) (Chen et al., 2018). Thus, the detection scheme of Cas12a is easier to design and simplifies the process, making it more suitable for POC diagnostics. Furthermore, PAM of Cas12a is T rich region and is different from Cas9 with G rich region, which provides a collaborative possibility of Cas9 and Cas12a (Yao et al., 2018). Overhangs of dsDNA by Cas12a cleavage combining with blunt ends cleaved by Cas9 could also support a large diversity of gene editing.

Unlike ELISA assays which are relied on delicate reagents, Cas12a assay is still active after 24 hrs, demonstrating high robustness, and is suitable for in-field diagnostics. Compared with PCR, which requires sample preparation, it has been shown that Cas12a assay can directly work with bodily fluids, thus avoiding complicated target extraction (Chen et al., 2018).

The hybridization of DNA-crRNA-LbCas12a complex is the key to activate the system and it shows excellent target specificity. For example, A-DNA and M-DNA have five different nucleobases out of 274 nt target. The mutation of the M-DNA prevents its binding with crRNA. Without crRNA binding, the CRISPR system cannot be activated for probe cleavage. More importantly, longer incubation time only increases the intensity of A-DNA but not M-DNA, thus achieving a better detection sensitivity. The high specificity of CRISPR Cas12a assay is poised to pair with other CRISPR Cas proteins such as Cas13a for multiplexing pathogen detection.

## 5. Conclusion

In this work, we present a complete and compact POC system for rapid and all-solution phase ASFV detection. It is based on a CRISPR Cas12a assay to trigger the indiscriminate ssDNA denaturation. The fluorescence signal is measured by a small and sensitive fluorescence sensing unit with a disposable cartridge. Without target amplification, a detection limit of ∼100 fM is achieved. This integrated system is ready to be used for POC detection of ASFV and other DNA based pathogens.

## 6. Acknowledgment

The authors would like to thank Prof. Mitchell R. O’Connell at the University of Rochester for valuable discussions. Qian He would like to acknowledge the graphic design by Qianhui Shi and research support by Prof. Julie A. Thomas at RIT. Ke Du would like to thank for the support by the Personalize Healthcare Technology Center (PHT180) at RIT.

